# Acid stress modulates metabolo-inflammatory pathways in oral epithelial cells

**DOI:** 10.64898/2026.03.16.711383

**Authors:** Angela Chen, Karen Zhu, Cameron T. Dixon, Adam D. Lietzan, Christina L. Graves

**Author notes:** **Correspondence** Christina L. Graves.

## Abstract

Acidification of the oral environment has been implicated in the initiation and progression of oral pathologies including oral cancer, but how acidic environments modulate normal oral epithelial cell (OEC) responses to microbial ligands is not understood. This study examined the impact of acidic stress on OEC morphological, molecular, and functional responses to toll-like-receptor ligand engagement *in vitro*. OEC cultures were exposed to either normal (pH:=:8.0) or acidified growth media (pH:=:3.0) for 24 hours prior to machine-learning-guided morphological analysis and exposure to either toll-like receptor (TLR)5 (flagellin) or TLR2/TLR1 (Pam3CSK4) agonists. Multiplex gene expression technology was used to quantify the transcriptional responses of metabolic-and immune-related genes at 6 hours post-TLR agonist exposure. OEC-mediated production of transforming growth factor-beta (TGF-β) was assessed by enzyme-linked immunosorbent assay at 2-, 6-, and 24-hours post-agonist exposure. Results showed that acid exposure induced significant changes to OEC morphology resembling epithelial-mesenchymal transition, the differential expression of *n*=197 metabolic-and *n*=43 immune-related genes and significantly increased OEC TGF-β1 production. The results demonstrate that acid stress skews normal OECs towards pro-inflammatory and pro-oncogenic phenotypes when faced with concomitant microbial ligand challenge and provide key molecular clues to OEC survival strategies with potential implications for elucidating the early molecular events in the development of epithelial dysplasia.

**Article Highlights:**

- Acute acid exposure reduces survival of OECs
- A subpopulation of OECs is resistant to acid-mediated cell loss and undergo morphometric changes consistent with epithelial-mesenchymal transition
- Concurrent acid stress and TLR stimulation modulates transcription of immune and metabolic genes in OECs
- Acid stress increases TGF-β1 protein production of OECs following TLR agonist stimulation

## 1 Introduction

The oral cavity is continuously exposed to stressors that impact the local microenvironment and influence salivary buffering capacity and mucosal health. Local pH disruption has been implicated in the development and progression of oral cancers (1). In addition to the Warburg effect, which involves increased fermentation of glucose by cancer cells thereby producing an acidic microenvironment, risk factors for oral cancer such as tobacco and alcohol consumption have been shown to promote oral acidification and are linked to oral cancer development (2–7). Therefore, pH disruptions in the oral cavity are hypothesized to have a bidirectional impact on cancer initiation and progression.

Oral epithelial cells (OECs) directly interface with the external environment and play a critical role in coordinating immune responses to maintain mucosal homeostasis and barrier immunity (8–10). OEC activation by bacterial virulence factors or local damage mediates innate immune cell recruitment and promotes inflammation and immunomodulatory responses (11). In oral cancer, dysregulated OEC Toll-like receptor (TLR)-mediated sensing of microbial byproducts has been shown to promote a proinflammatory phenotype that contributes to a pro-oncogenic and immunosuppressive microenvironment in oral cancer (12–14). Studies suggest that acid exposure has been shown to modulate epithelial immune responses *in vitro* and regulate epithelial-mesenchymal transition (15, 16). However, little is known about how acid impacts the immunological and metabolic properties of healthy oral epithelium. In this study, we examined the effects of acid on OECs under TLR agonist simulation, providing insights into the molecular mechanisms mediating OEC adaptation to acidic stress in the context of microbial ligands.

## 2 Material and Methods

### 2.1 Cell culture

Low-passage mixed-donor, Primary Human Oral Epithelial Cells (Cat# 36063-01, Celprogen, Inc., Torrance, USA), were thawed from liquid nitrogen, plated on poly-L-lysine-coated T-75 cell culture flasks (Sigma-Aldrich Co., St. Louis, USA; Corning Inc., Durham, USA) and maintained in Human OEC Culture Complete Growth Media (Celprogen, Inc.) containing serum and antibiotics at 37°C and 5% CO_2_ for two passages prior to experimentation according to manufacturer recommendations. Cells were seeded at a density of 0.05×10^6^ cells/well and at 80% confluence, cell cultures were acidified by addition of 1.0 N hydrochloric acid to the growth media (pH:=:3.0) (Sigma-Aldrich Co.). After 24h, standard culture media was replenished (pH:=:8.0) and cells were stimulated with flagellin (100 ng/mL) or Pam3CSK4 (1 µg/mL) (InvivoGen, San Diego, USA) for 2-, 6-, or 24-h. TLR-stimulated OEC in standard media (pH:=:8.0) were used as the experimental comparison. Each condition included n=4 replicates.

### 2.2 Microscopy and Image Analysis

Brightfield images were acquired on an EVOS^TM^ XL Core Imaging System fitted with a LPlan PH2 20x/0.40,∞/1.2 dry objective (Thermo Fisher Scientific Inc., Waltham, MA, USA). Brightness and contrast were linearly and uniformly applied to all images and converted to 16-bit grayscale prior to downstream analysis using FIJI (V2.14.0/1.54) software (17). Resulting micrographs were analyzed according to a custom computational pipeline using CellProfiler4 software (V4.2.6), CellPose as the base segmentation algorithm (18), and the RunCellpose, IdentifyPrimaryObjects, and MeasureObjectSizeShape modules. RunCellpose was used to generate cellular probability masks with a minimum size of 15 pixels (px), 20px expected object diameter, *n=*-6.0 cell probability threshold, and *n=*0.7 flow threshold. IdentifyPrimaryObjects was used to segment probability masks with a 20-85px diameter range using the Adaptive threshold strategy and the Sauvola thresholding method. Window size was adjusted to 20-25px with a *n=*0 smoothing scale, *n*=15-30 smoothing filter, and *n=*0.9 threshold correction factor. Lower and upper threshold bounds were fine-tuned between *n=*0.1 and-1.0. Intensity was used to demarcate dividing lines between clumped objects with a 7-15px local maxima suppression.

Cell count, area, perimeter, and form factor were obtained via segmentation processing using MeasureObjectSizeShape.

### 2.3 Molecular Profiling

Total RNA was harvested using a RNeasy Mini Kit according to manufacturer instructions (Qiagen Inc.). RNA purity and quantity were measured using a NanoDrop™ One Microvolume UV-Vis Spectrophotometer (Thermo Fisher Scientific Inc.). Changes in OEC transcriptional activity after 6h TLR stimulation were measured using the nCounter® XT_HuV2_Immunology RLF Innate Immune Profiling and nCounter® XT_Hs_Metabolic_Path RLF Metabolic Profiling code set panels according to manufacturer supplied protocols (NanoString Technologies Inc., Seattle, USA). Hybridization was performed using 100ng of RNA per sample for 18h at 65°C. Data were analyzed by ROSALIND® (https://rosalind.bio/), with a HyperScale architecture developed by ROSALIND, Inc. (San Diego, CA, USA). Fold changes and *p*-values were calculated using the fast method as described in the nCounter® Advanced Analysis 2.0 User Manual. *P-*value adjustment was performed using the Benjamini-Hochberg method of estimating false discovery rates (FDR). Read Distribution percentages, violin plots, identity heatmaps, and sample MDS plots were generated as part of the QC step. Normalization, fold changes, and p-values were calculated using criteria provided by NanoString. ROSALIND® follows the nCounter® Advanced Analysis protocol of dividing counts within a lane by the geometric mean of the normalizer probes from the same lane. Housekeeping probes to be used for normalization are selected based on the geoNorm algorithm as implemented in the NormqPCR R library. Abundance of various cell populations is calculated on ROSALIND® using the Nanostring Cell Type Profiling Module. ROSALIND® performs a filtering of Cell Type Profiling results to include results that have scores with fold changes by ±1.25 and *p*≥0.05. Data are publicly available at github.com/cgraveslab/OEC-stress.

### 2.4 ELISA

Concentration of TGF-β1 in OEC culture supernatants was determined using a Human ELISA Kit (Invitrogen, Cat#: BMS249-4/TEN) according to manufacturer’s instructions and loaded onto assay plates in technical duplicate for each biological replicate. The optical density (O.D.) was read at 450 nm with a 620 nm pathlength correction on an Epoch colorimetric plate reader using Gen5(V2.0.5) Software (BioTek, Winooski, VT, USA). Background-corrected O.D. values were transformed to concentration (pg/mL) using an 8-point standard curve and normalized via log-transformation to enable statistical comparison.

### 2.5 Statistics

GraphPad Prism V10.4.1 (GraphPad Software Inc., La Jolla, CA, USA) was used for graphical representation and statistical analysis unless otherwise noted. Each dataset was tested for normality and lognormality prior to statistical comparison. T-test or ANOVA were used for two-or more-group comparisons as appropriate and as detailed in each corresponding figure legend.

## 3 Results

### 3.1. Acid stress reduces cell number and disrupts oral epithelial cell morphology

In health, salivary pH ranges from 6.2 – 7.6 with variables such as diet and reduced salivary buffering capacity having the ability to drive the pH <5.0 (19–21). Previous studies have shown that acidification of the microenvironment can regulate epithelial cell morphology and function including epithelial-mesenchymal transition (15). Additionally, although acidification is an important component of the tumor microenvironment, how acid modulates OEC morphology, gene transcription, and function in is largely unknown (1).

To investigate the impact of acid on OECs, we cultured OECs in media buffered to pH==3.0 and performed morphometric analysis prior to downstream TLR stimulation, molecular profiling, and functional analysis (**Fig 1**). Brightfield micrographs were obtained and evaluated using an open source, machine-learning based analysis pipeline (**Fig 2A-C**). As expected, acid resulted in a significant reduction in cell number and total area coverage suggesting that acid causes significant *in vitro* cell loss (**Fig 2D-E**). Additionally, cell tracing found that acid challenged OECs exhibited larger morphological variety as evidenced by significant increases in cellular area, perimeter, and form factor (**Fig 2F-H**).

**Figure 1.**
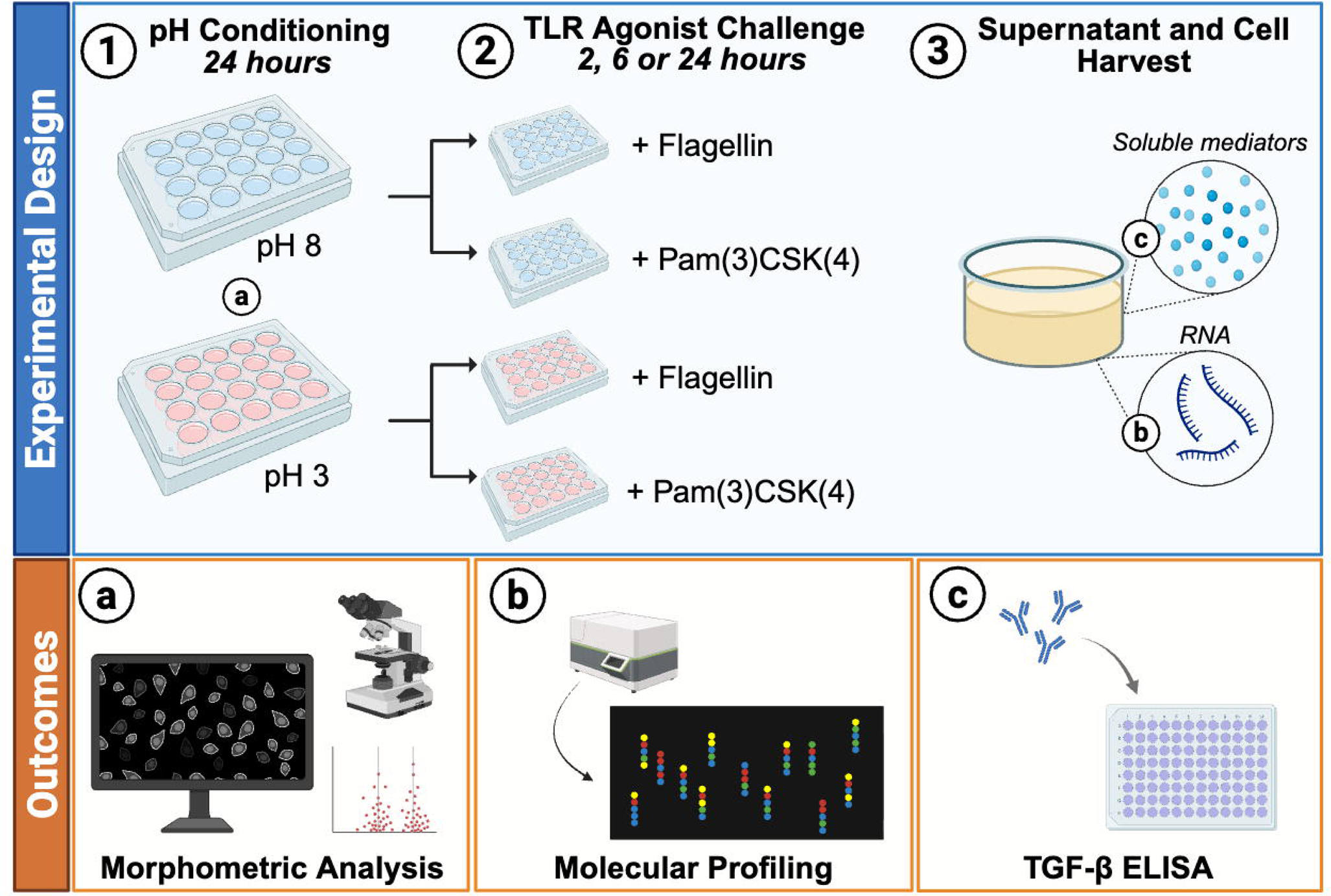
Experimental Overview. Schematic representation of the experimental design to investigate the impact of acid stress on oral epithelial cells (OECs) and Toll-like receptor (TLR) agonist stimulation. **pH Conditioning** (**1**): OECs (80% confluence) were cultured in acidified (pH:=:3.0) or complete growth media (pH:=:8.0) for 24h. **Morphometric Analysis** (**a**): Brightfield micrographs and a machine-learning based image analysis pipeline were used to assess changes in cellular morphology. **TLR Agonist Challenge** (**2**): Cell cultures were subjected to either 100 ng/mL flagellin (TLR5 agonist) or 1 mg/mL Pam3CSK4 (TLR2/1 agonist) for 2-, 6-, or 24h. **Molecular profiling** (**b**): was conducted on OEC RNA collected after 6h TLR agonist challenge using NanoString® nCounter® technology followed by pathway analysis using the Gene Ontology knowledgebase. A **TGF-**β **ELISA** (**c**) was performed on OEC supernatants collected 2-, 6-, and 24h post TLR agonist stimulation. Figure made with Biorender.com

**Figure 2.**
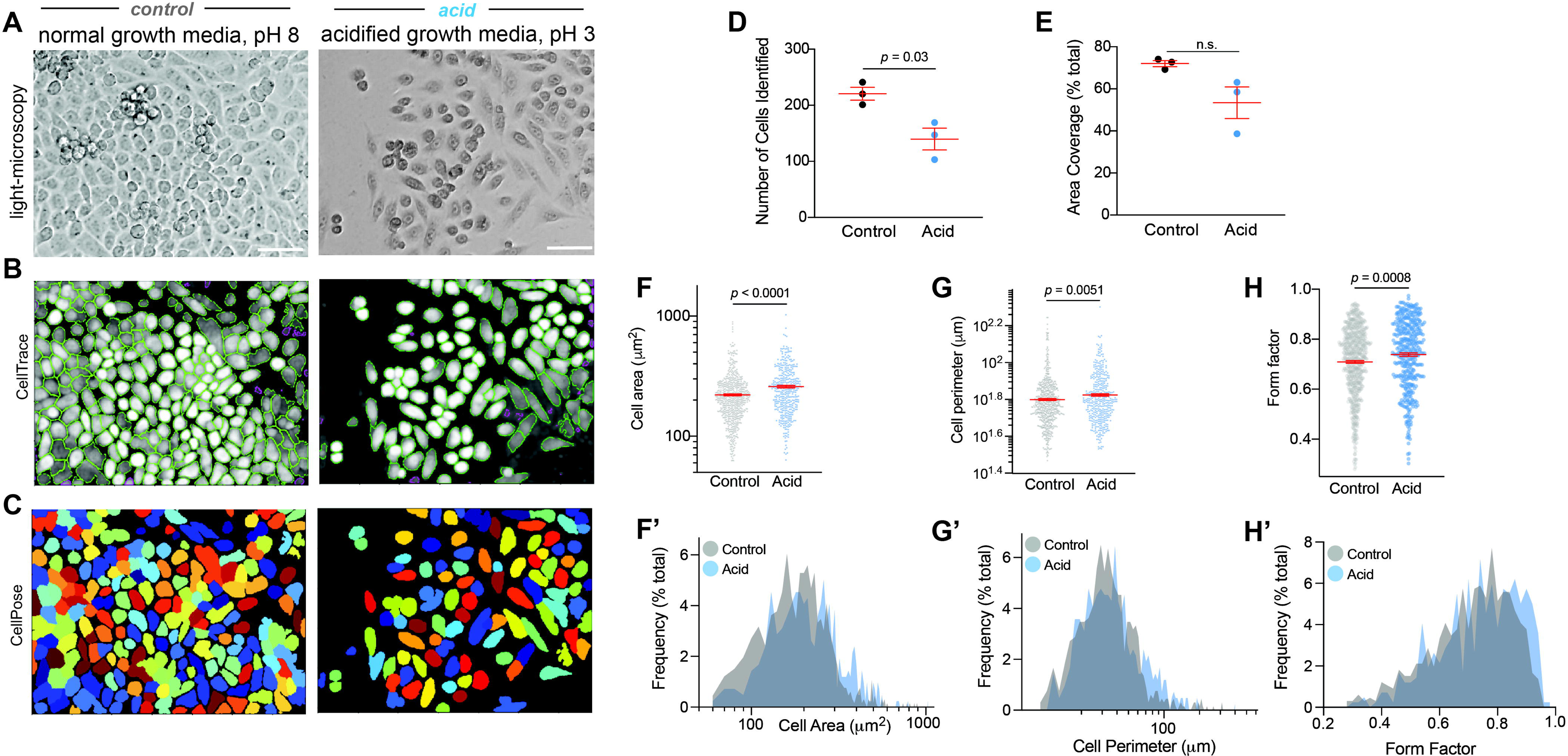
Acid stress alters OEC morphology. OEC cultured in the presence of pH:=:8.0 (control, left) or pH:=:3.0 (experimental, right) growth media prior to TLR agonist stimulation shown as (**A**) brightfield micrographs, (**B**) CellTrace masks or (**C**) CellPose masks. (**A-C**) Images shown are representative of n=3-4 images and n=24 observations per condition. Scale bar = 50 μm. CellTrace masks were used to quantify OEC (**D**) number and (**E**) area coverage. Representative dot plots depicting (**F**) cellular area (mm^2^), (**G**) cellular perimeter (mm), or (**H**) form factor on a per-cell basis. Corresponding frequency histograms (% of total objects identified) for **F, G,** and **H** are shown in **F’**, **G’**, and **H’**, respectively; each dot represents one cell mask. (**D-E**) Error bars show mean ± S.D. *p*-values derived from Welch’s T-test are shown for each comparison. n.s.=not significant. Blue dots/bars=acid; black and grey dots/bars=control. Data are representative of n=3 independent runs and n=2-3 image fields.

The oral epithelium is continually exposed to microbes and their byproducts, and this interaction persists irrespective of salivary pH (22). To probe the impact that acid has on OEC microbial recognition and responsiveness, flagellin or Pam3CSK4, which mimic dominant microbially associated proteins found in plaque biofilm, was introduced to the cultures. After stimulation, total RNA was harvested and quantified (**Fig 1, Supplementary Fig 1**). Consistent with our morphometric findings, acid-stressed OEC cultures exhibited a 4-fold reduction in recovered RNA, independent of agonist (**Supplementary Fig 1A**). Acid stress induced RNA absorbance profile shifts including a slight-but-significant reduction in the A260/280 ratio (**Supplementary Fig 1B**) and significant reduction in the A260/230 ratio (**Supplementary Fig 1C**) suggesting elevated apoptotic activity, accumulation of oxidized molecular byproducts, and/or chromatin remodeling.

### 3.2. Engagement of immune pathways following acid stress

Work from our group and others has previously demonstrated that epithelial cells, including OECs, can function as immune mediators in health and disease (23–25). We therefore hypothesized that when faced with an acidic environment prior to TLR stimulation, OECs might differentially engage immune-mediated pathways. To test this, we conducted a multiplexed gene expression analysis of immune-related genes (**Fig 1**). We found that acid stress was associated with the differential expression of 24 genes (19 upregulated; 5 downregulated) during flagellin stimulation (**Fig 3A-C**) and 19 genes when stimulated with Pam3CSK4 (16 upregulated; 3 downregulated) (**Fig 3D-F**). Evaluating immunological gene expression similarities between the TLR agonists, a modest 12 genes were shared between the agonists and significantly upregulated (**Supplementary Table 1**). These genes were broadly involved with signaling pathways for NF-κB (RELA, MALT1, NFKBIZ, and CHUK), MAPK (MAP4K4, HRAS), transcriptional regulation (SKI, STAT6, NFATC3, ZEB1, and TCF4), and proteosome function (PSMD7). Altogether these results suggest that acid and TLR stimulation promote OEC plasticity and transcriptional reprogramming of genes involved with immune priming, proteolytic processing, and transcriptional regulation.

**Figure 3.**
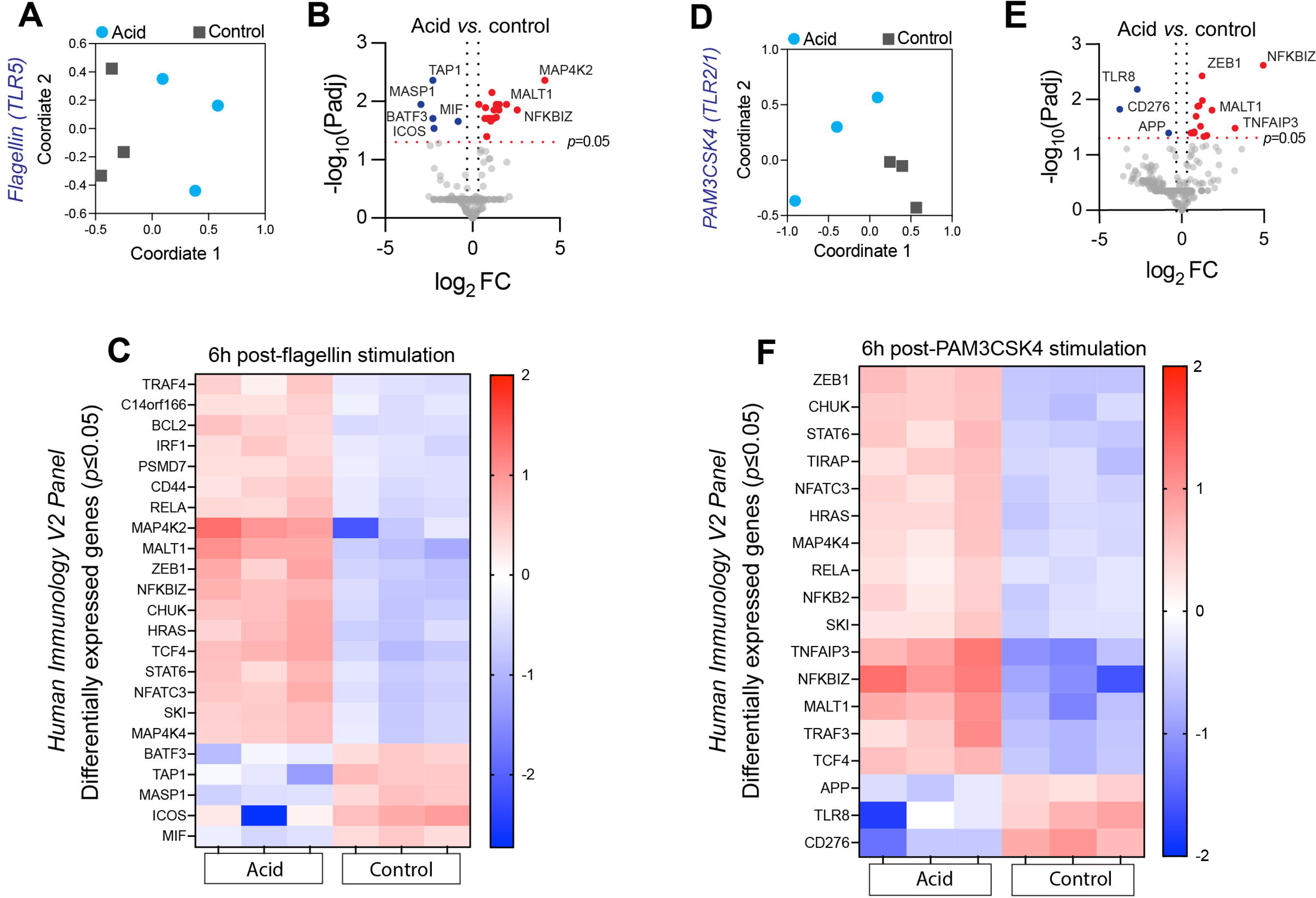
**Acid stress modulates OEC immune responses to TLR agonists.** MDS plots (**A,D**), volcano plots (**B, E**) and heatmaps (**C, F**) showing differential immune-related gene expression in OECs after exposure to pH:=:3.0 or pH:=:8.0 growth media and subsequent flagellin (100 ng/mL) or Pam3CSK4 (1µg/mL) agonist stimulation (6 hours), respectively. (**A,D**) blue dots=acid; grey squares=control; (**B, E**) Blue dots=*p*<0.05, FC<-1.25; Red dots=*p*<0.05, FC≥1.25. Grey dots=n.s. Heatmaps show only significantly (*p*≤0.05) differentially regulated genes.

When focusing on molecular profiles unique to each agonist, only 4 genes were upregulated and unique to acid stress in the context of Pam3CSK4 stimulation. These genes included NFKB2, TIRAP, TRAF3, and TNFAIP3 (A20), which are classically associated with immune modulation suggesting that acid enhances canonical and non-canonical NF-κB responses in OECs under TLR2/1 agonist stimulation. Conversely, acid in the context of flagellin stimulation promoted an increase in gene expression associated with TGF-β signaling and epithelial-mesenchymal transition (EMT) including TGF-β Receptor 2 (TGFBR2), TNF Receptor Associated Factor 4 (TRAF4), and CD44 Antigen (CD44); immune and inflammatory signaling including Interferon Regulatory Factor 1 (IRF1) and TRAF4; regulation of apoptosis (BCL2, and IRF1), the MAPK/JNK Stress Response Pathway (MAP4K2); and RNA processing and translation, indicated by upregulation of CD14orf166. qPCR analysis of acid-stressed cultures demonstrated a significant increase in transcription of vimentin, a marker of EMT (**Supplementary Figure 2**). These results indicate that acid stress enhances flagellin-mediated TGF-β signaling and EMT responses in OECs and suggests that concomitant acid stress and NF-κB activation may sensitize OECs to subsequent TLR stimulation.

Interestingly, no downregulated immune-related genes were shared between the flagellin and Pam3CSK4 stimulation conditions suggesting that OEC responses to TLR agonists are context-dependent. We found 3 genes to be significantly downregulated due to acid unique to Pam3CSK4 stimulation, including Toll-like Receptor 8 (TLR8), Cluster of Differentiation 276 (CD276/B7-H3) and Amyloid Precursor Protein (APP) suggesting that acid stress impairs homeostatic OEC immunological responses to Pam3CSK4. Conversely, 5 genes were downregulated due to acid that were unique to flagellin stimulation including Macrophage (mφ) Migration Inhibitory Factor (MIF), Inducible T-cell Co-Stimulator (ICOS), Transporter Associated with Antigen Processing 1 (TAP1), Basic Leucine Zipper ATF-Like Transcription Factor 3 (BATF3), and Mannan-Binding Lectin Serine Protease 1 (MASP1). Altogether these results show that acid and concomitant TLR agonist stimulation results in divergent downregulation of immune-related genes in OECs.

### 3.3. Acid stress induces metabolic reprogramming of OECs in the context of TLR agonist stimulation

We next hypothesized that acid stress would have a marked effect on metabolism-related gene expression, particularly given our observations that acid induced remarkable changes to cellular morphology and ribonucleic acid absorbance profiles. Here, acid stress prior to Pam3CSK4 stimulation resulted in the differential expression of 116 metabolic genes (104 upregulated; 12 downregulated) while acid prior to flagellin stimulation resulted in differential expression of 81 metabolic genes (44 upregulated, 37 downregulated) (**Fig 4, Supplementary Table 1**). Overall, metabolism-related genes were mostly upregulated and largely shared in response to acid and either subsequent flagellin or Pam3CSK4 stimulation although Pam3CSK4 appeared to have a stronger effect. Specifically, 37 genes were upregulated due to acid in response to either flagellin or Pam3CSK stimulation.

**Figure 4.**
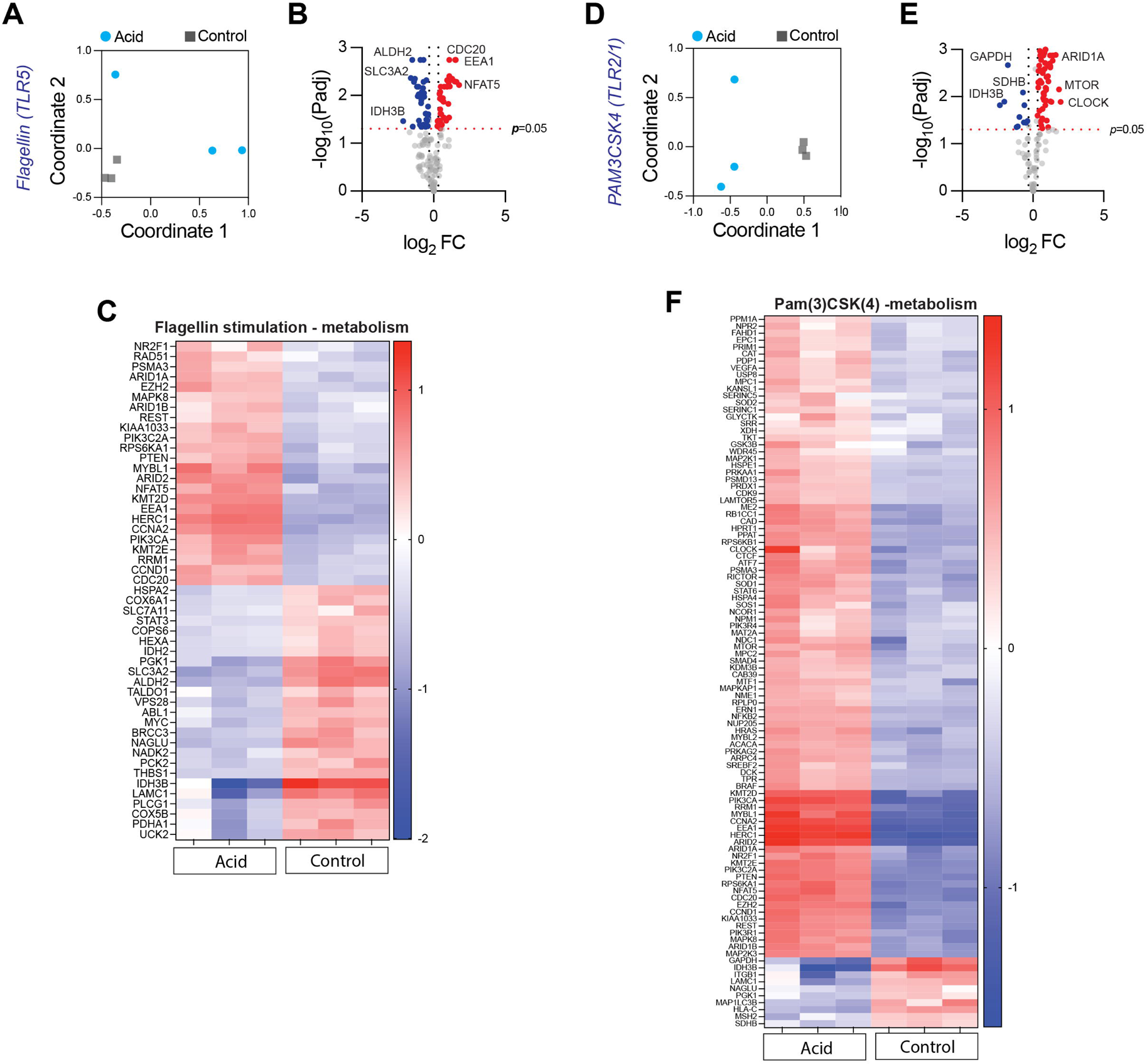
Acid stress modulates OEC metabolic responses to TLR stimulation *in vitro.* MDS plots (**A, D**), volcano plots (**B, E**) and heatmaps (**C, F**) showing differential metabolic OEC gene expression after exposure to acidified (pH:=:3) or control (pH:=:8) growth media and subsequent flagellin (100 ng/mL; shown in **A-C**) or Pam3CSK4 (1 µg/mL; shown in **D-F**) agonist stimulation (6 hours), respectively. (**A, D**) blue dots=acid; grey squares=control; (**B, E**) Blue dots=*p*<0.05, FC<-1.25; Red dots=*p*<0.05, FC>1.25. Grey dots=n.s. Heatmaps show only significantly (*p*≤0.05) differentially regulated genes.

Intriguingly, acid stress resulted in the upregulation of only one gene, RAD51 – a gene involved in the repair of double-stranded DNA breaks, critical for protecting genome integrity – that was specific to flagellin stimulation. In response to Pam3CSK4, acid stress resulted in the engagement of genes associated with cell cycle progression, chromatin remodeling, DNA synthesis and repair, vesicle trafficking and signal transduction (**Fig 4, Supplemental Table 1**). Intriguingly, acid stress resulted in a Pam3CSK4-specific upregulation of CLOCK, a key orchestrator of circadian regulation, genes associated with angiogenesis including VEGFA, and lipid metabolism (SERIN). Among all differentially regulated genes, the most dramatically upregulated genes in response to acid post-Pam3CSK4 were KMT2D, PIK3CA, and RRM1, which were also engaged due to acid post flagellin stimulation. Overall, these results suggest that acid stress may result in a hyperproliferative, potentially oncogenic or regenerative state in OECs.

Acid stress caused an upregulation of genes associated with proliferation and cell cycle acceleration including PI3K/AKT/mTOR pathway engagement (growth and survival), and engagement of Cyclin A2 (CCNA2) and Myb proto-oncogene like 1 (MYBL1). Taken together these data show that acid-stressed cells are primed for cell division, DNA replication, and resistance to apoptosis. Dramatic upregulation of genes associated with chromatin remodeling & transcriptional activation (KMT2D, ARID2) suggest a global loosening of chromatin and broad transcriptional activation in response to acidic stress. Genes involved in intracellular trafficking and signal modification (HECT, HERC1) and Early endosome Antigen 1 (EEA1) were also upreglated, indicative of increased membrane turnover and receptor recycling.

Several metabolic genes were downregulated due to acid unique to flagellin stimulation (**Supplementary Table 1**, **Fig 4**) including genes related to cellular metabolism, including PCK2, PDHA1, PFKM, and ADK), mitochondrial function (COX5B, COX6A1, and COX4I1), solute transport (SLC7A5 and SLC7A11), and protein degradation (VPS28, COPS6, PSMC1). Intriguingly, a downregulation of known tumor proto-oncogenes including AKT2, MYC, ABL1, STAT3, and KRAS was also observed. Acid stress unique to Pam3CSK4 stimulation resulted in significant downregulation of 6 genes including SDHB, HLA-C, GAPDH, MSH2, ITGB1, and MAP1LC3B indicating that acid stress impairs cellular metabolism, reduces immune surveillance, DNA repair mechanisms and autophagy. In contrast to uniquely engaged gene networks, acid stress downregulated key cellular energy networks, irrespective of subsequent TLR agonist stimulation. These include engagement of the lysosomal enzymes and basement membrane components (NAGLU, LAMC1), metabolic enzymes (IDH3B, PGK1, ALDH2), and solute carriers (SLC3A2/CD98hc) suggesting that acid stress downregulates key cellular energy networks.

### 3.4. Acid stress induces TGF-β1 production in oral epithelial cells

Given that multiple differentially regulated genes are known to be upstream of TGF-β1 (Fig 3, Fig 4), we assessed acid-mediated OEC production of TGF-β1 following TLR agonist stimulation. Acid stress temporally increased TGF-β1 production, with a significant increase observed at 24h post-stimulation for both agonists (**Fig 5**). Given that similar levels of TGF-β1 were recovered under either stimulation condition, these results suggest that acid stress primes OECs for heightened TGF-β1 production following microbial ligand engagement.

**Figure 5.**
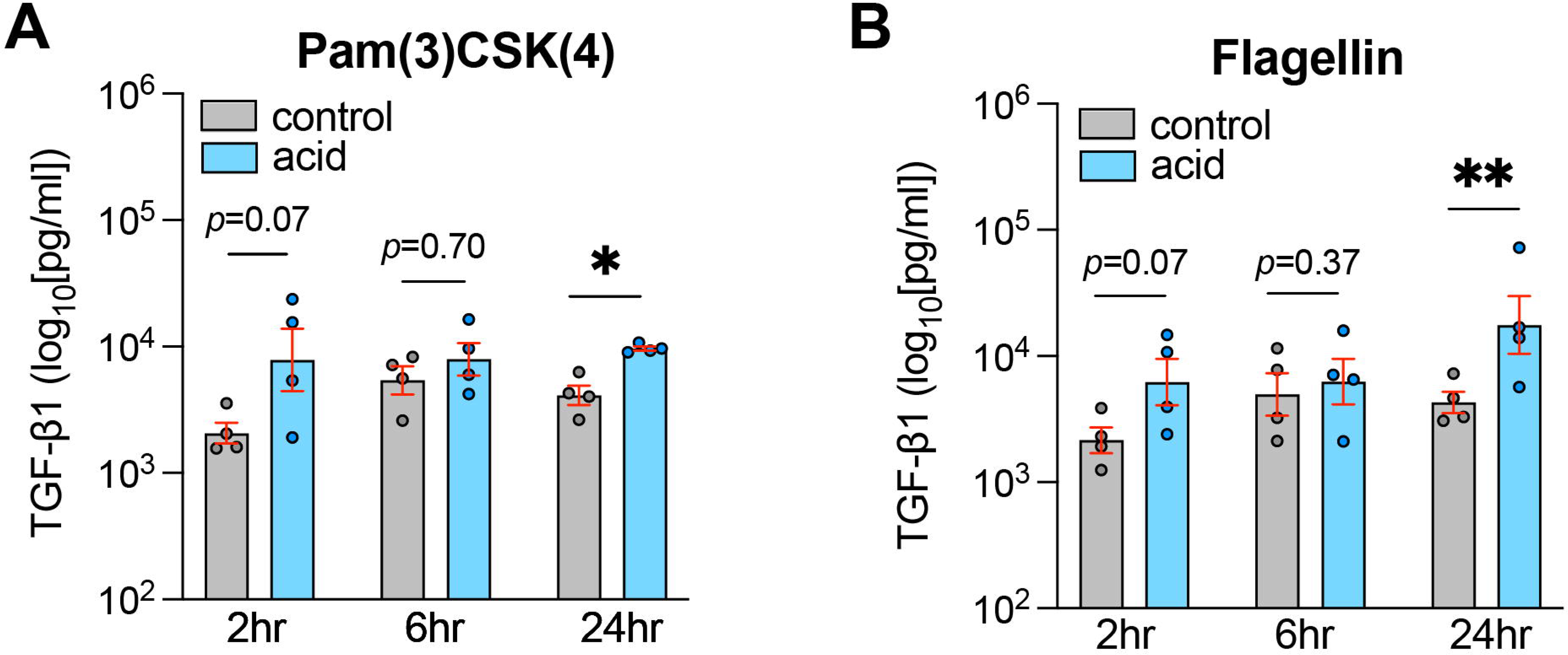
Acid stress increases OEC TGF-b1 production. TGF-β1 protein concentration produced from OEC cultures after a 24h acid exposure and following 2-, 6-, or 24-hours of (**A**) Pam3CSK4 or (**B**) Flagellin stimulation. Grey bars=supernatant from control media cultures. Blue bars = supernatant from acid exposed cultures. Resulting values (pg/mL) were normalized to total RNA recovered in each well and log-transformed to enable statistical comparison by Student’s T-test. \**p*<0.05; \*\**p*<0.01.

## 4 Discussion

Acidic oral microenvironments can be caused by a wide variety of salivary, dietary, microbial, host, physiological, medical, pharmacological, behavioral and environmental factors including hypoxia, inflammation, acid reflux, or poor oral care (26–29). In addition, epidemiological studies have demonstrated that dietary acid load is associated with both increased risk as well as poor prognosis in cancer including oral cancer (30, 31). Despite these associations, the influence of acidic microenvironments on healthy oral epithelia is largely unknown and the mechanisms underlying adaptive OEC responses to acidic stress are incompletely understood.

In this study, we assessed the impact of acid stress on OEC morphology and microbial ligand responsiveness (**Fig 1**). Acid stress resulted in a significant reduction of OECs, likely owing to apoptotic, necrotic or anoikis-related cellular death (**Fig 2, Supplementary Fig 1**). Interestingly, despite a significant reduction in cell number, remaining OECs exhibited profound morphological changes consistent with EMT including cellular elongation and increased size (**Fig 2**).

Prior work has shown that epithelial cells, including OECs, modulate mucosal homeostasis in part through TLR engagement, and play a critical role in both inflammatory responses and immunoregulation (23, 25, 32, 33). In addition, NF-κB pathway activation has been shown to promote inflammation, cellular survival and proliferation in oral cancer (11). In this study, OEC morphological changes in response to acid stress were observed alongside profound transcriptional changes to immune and metabolism-related gene expression signatures in the context of microbial ligand stimulation.

In response to acid following flagellin stimulation, acid induced gene signatures consistent with stemness, EMT & migration (TGFBR2, TRAF4, CD44) and regulation of apoptosis (BCL2) were observed altogether suggesting that acid stress selects for a subpopulation of OECs capable of withstanding acidic environments and subverting cell death through EMT (**Fig 2-3**, **Supplementary Table 1**). In response to acid following Pam3CSK stimulation and irrespective of agonist, genes involved in NF-κB pathway activation, MAPK and Ras signaling, and T cell activation were upregulated, consistent with immune activation and EMT (**Fig 2-3**, **Supplementary Table 1**). While HRAS overexpression is well-appreciated to drive cellular proliferation and tumorigenesis, we also observed upregulation of genes related to transcriptional reprogramming and EMT including ZEB1 and TCF4 which have been demonstrated to enhance OEC invasiveness and metastatic potential (34, 35).

Our data also showed that acidic stress primes OECs to dampen microbial clearance, mφ-T cell interactions, and homeostatic T cell functions in response to flagellin including downregulation of macrophage migration (MIF), T-cell co-stimulation (ICOS), antigen processing (TAP1, BATF3), and complement responses (MASP1) (**Fig 3, Supplementary Table 1**). Taken together these results suggest that acid stress may be an important environmental factor that dysregulates OEC-mediated mφ and T cell responses and impairs homeostatic immunological responses to flagellin.

Acid-dependent suppression of TLR8 responses post-Pam3CSK4 stimulation was also observed suggesting that acidic stress could impair responses to single-stranded RNA viruses potentially resulting in impaired oral antiviral responses. Downregulation of immune checkpoint inhibitors including CD276/B7-H3 suggest that acid-stressed OECs could be more susceptible to T/NK cell killing but might also allow for uncontrolled inflammation since T/NK cell activation is critical for T cell suppression in the mucosa. We also observed a downregulation of Amyloid Precursor Protein (APP) in acid-stressed OECs following Pam3CSK4 stimulation. In healthy epithelium, APP is broadly expressed at low levels where studies show it regulates the secretion of amyloid peptides for barrier function, regulates cholesterol uptake, promotes inflammatory responses, and exerts antimicrobial activity. Emerging studies show that APP is overexpressed in several cancers including pancreatic, lung, colon and prostate cancer but whether APP could be an oral cancer biomarker is unknown (36).

Acid and TLR-agonist stimulation resulted in the differential expression of n=197 metabolic genes. Interestingly, only one gene – RAD51 – was upregulated in response to acid specific to flagellin stimulation. Overexpression of RAD51 has been observed in various cancers and has been suggested to be a prognostic factor in oral cancer demonstrating that RAD51 may enable OEC persistence during oxidative stress, similar to its homologous function in the oral pathogens *Streptococcus mutans* and *Porphyromonas gingivalis* (37–40). Our data also revealed that acid stress upregulated circadian rhythm-associated genes including CLOCK in the presence of Pam3CSK4 (**Fig 4E, F**). Interestingly, macrophage circadian rhythm has been shown to be disrupted by acidic tumor microenvironments (41). Acid stress also resulted in SLC3A2 overexpression in OECs. Interestingly, SLC3A2 has been observed in oral cancer where overexpression has been shown to be involved in increased migration, invasion, and proliferation of cancer cells with key roles in regulating apoptosis (42, 43).

Acid significantly increased OEC TGF-β1 production following TLR stimulation (**Fig 5**). TGF-β1 has also been shown to play a pivotal role in cancer development including oral squamous cell carcinoma by facilitating tumor development, evading immune surveillance, and EMT through metabolic reprogramming of amino acid metabolism (44–49). Our data suggest that OEC responses to acidic stress may be an underappreciated factor in TGF-β1 dysregulation where increased TGF-β1 links immunological and metabolic function of OECs. Taken together, these results indicate that acidic microenvironments may exacerbate TGF-β1 production, providing a potential mechanism linking acid stress to the development or progression of epithelial-driven oral diseases associated with malignancy, such as oral lichen planus (50–52).

### 4.1 Limitations & Future Directions

In this study, we considered OEC responses under very low pH (pH=3.0) exposure, which may be most relevant to gastroesophageal reflux or chronic sweetened beverage exposure; future work will explore milder acidification conditions. Second, pH and microbial ligand stimulation was performed for relatively shorter periods of time; we are therefore unable to comment on persistent or repeated effects of low pH or microbial stimulation on OECs. Future studies should explore whether OECs can adapt to a low pH environment, as these studies could reveal molecular clues as to how chronic inflammation drives early oncogenic processes.

### 4.2 Conclusions

Altogether our findings demonstrate that acidic stress in the context of microbial ligand stimulation contributes to pro-inflammatory and pro-oncogenic phenotypes in OECs by disrupting cellular morphology consistent with anoikis and EMT, modulating immune and metabolic gene expression and promoting TGF-β1 secretion. These data suggest that oral acidification may play a key role in early immune-and metabolic-dysregulation consistent with OEC dysplasia, transcriptional activation, and epigenetic remodeling potentially underlying early malignant transformation events.

## 6 Author Contributions

A.C.-conception and design, data acquisition, analysis, and drafting of the manuscript. K.Z.-data acquisition, analysis and drafting of the manuscript. C.D.-data acquisition, analysis, drafting, and manuscript revision. A.L.-data interpretation, drafting, and manuscript revision. C.G.-conception, design, data acquisition, analysis, interpretation, drafting and manuscript revision.

All authors gave final approval and agreed to be accountable for all aspects of the work ensuring integrity and accuracy.

## Supporting Information

**Supplementary Figure 1. Acid stress effects on RNA recovery and absorbance.**

**(A)** Total RNA concentration (ng/mL) recovered from control (grey bars) or acid-stressed cells (blue bars) following Pam3CSK4 (left) or flagellin (right) stimulation. (**B, B’**) A260/280 and (**C**) A260/230 ratio of purified RNA. For panels **B** and **C**, all control (n=24) or acid (n=24) samples were pooled for statistical comparison (Mann-Whitney test).

**Supplementary Table 1. Differentially expressed gene lists (padj<0.05, |FC|>1.25).**

Metabolic and Immune-related genes are categorized based on up-or down-regulation and further grouped TLR agonist. Genes that appear up-or down-regulated for TLR agonists are listed in under “Shared”. *Indicates gene appears in both the metabolic and immune gene panels.

**Supplementary Figure 2. Acid stress induces transcriptional upregulation of vimentin.**

qRT-PCR data showing the relative expression of vimentin, a marker of EMT, in OEC after 24h exposure to a low pHe environment. Each dot represents one biological replicate. All control (n=24) or acid (n=24) samples were pooled for statistical comparison (Mann-Whitney test).

