## Supplemental Information for "Acid stress modulates metabolo-inflammatory pathways in oral epithelial cells"

**Supplementary Figures**

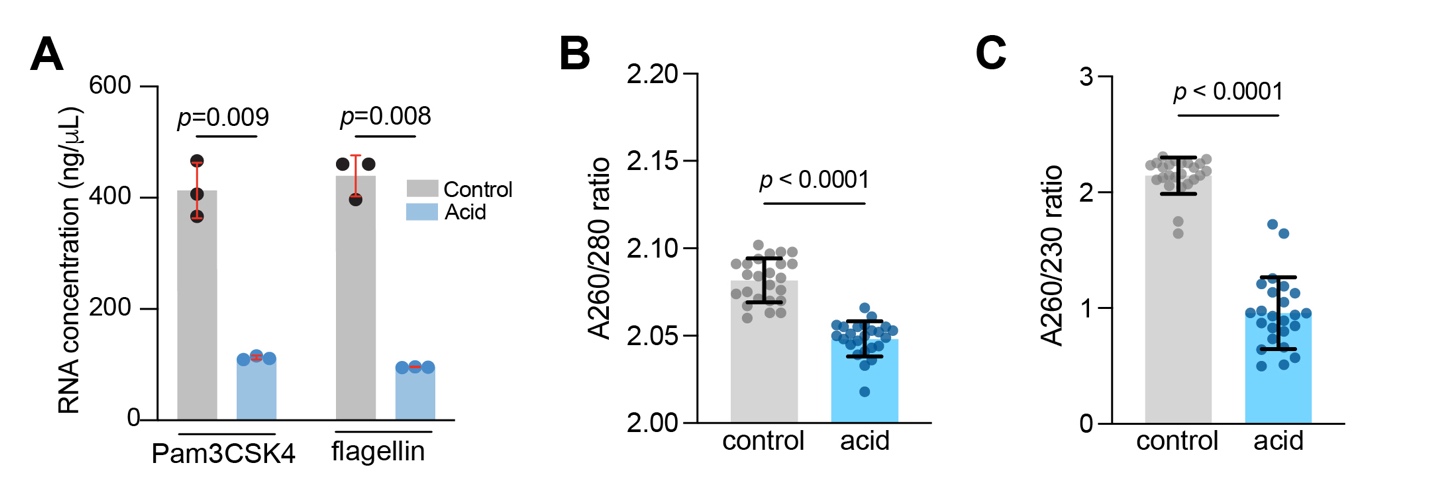

### **Supplementary Figure 1. Acid stress effects on RNA recovery and absorbance.**

| **Upregulated Immune Genes** | | | | | | | |
| --- | --- | --- | --- | --- | --- | --- | --- |
| **Flagellin** | TGFBR2 | C14orf166 | TRAF4 | CD44 | IRF1 | MAP4K2 | BCL2 |
| **Shared** | PSMD7 | RELA | MAP4K4 | SKI | STAT6 | MALT1 | NFKBIZ |
|  | NFATC3 | CHUK | HRAS* | ZEB1 | TCF4 |  |  |
| **Pam3CSK4** | NFKB2* | TIRAP | TRAF3 | TNFAIP3 |  |  |  |
| **Downregulated Immune Genes** | | | | | | | |
| **Flagellin** | MIF | ICOS | TAP1 | BATF3 | MASP1 |  |  |
| **Shared** |  |  |  |  | |  |  |
| **Pam3CSK4** | APP | TLR8 | CD276 |  | |  |  |
| **Upregulated Metabolic Genes** | | | | | | | |
| **Flagellin** | RAD51 |  |  |  |  |  |  |
| **Shared** | NFAT5 | HERC1 | KMT2D | MYBL1 | EEA1 | MYBL2 | HPRT1 |
|  | CCNA2 | ARID2 | PIK3CA | KMT2E | CDC20 | NUP205 | MTF1 |
|  | EZH2 | RRM1 | RPS6KA1 | ARID1A | CCND1 | ME2 | SOD1 |
|  | PTEN | PIK3C2A | KIAA1033 | NR2F1 | PSMA3 | DCK | CAD |
|  | ARID1B | MAPK8 | REST | STAT6 | TPR | RPS6KB1 | PPAT |
|  | MAP2K3 | ATF7 | PIK3R1 | BRAF | HRAS* | ARPC4 | HSPA4 |
|  | ERN1 |  |  |  |  |  |  |
| **Pam3CSK4** | CLOCK | MTOR | RICTOR | RB1CC1 | NDC1 | ARF5 | PFKM |
|  | SOS1 | CTCF | PRKAG2 | MPC2 | ACACA | ADK | SRR |
|  | NFKB2* | PDP1 | GSK3B | SREBF2 | GLYCTK | PSMC1 | WDR45 |
|  | SMAD4 | PRKAA1 | KDM3B | RPLP0 | NCOR1 | NDUFB7 | PPM1A |
|  | PIK3R4 | MAT2A | MAPKAP1 | CDK9 | NPM1 | SLC7A5 | XDH |
|  | NME1 | CAT | PSMD13 | LAMTOR5 | CAB39 | MGST3 | AKT2 |
|  | PRDX1 | HSPE1 | VEGFA | USP8 | MPC1 | LAMTOR2 | COX5A |
|  | EPC1 | SOD2 | MAP2K1 | FAHD1 | KANSL1 | GABARAP |  |
|  | PRIM1 | SERINC1 | TKT | SERINC5 | NPR2 | HSD17B8 |  |
| **Downregulated Metabolic Genes** | | | | | | | |
| **Flagellin** | BRCC3 | PLCG1 | VPS28 | PCK2 | PDHA1 | GABARAP | AKT2 |
|  | MYC | UCK2 | THBS1 | ABL1 | COX5B | SLC7A5 |  |
|  | NADK2 | HSPA2 | COX6A1 | IDH2 | TALDO1 | NDUFB7 |  |
|  | STAT3 | COPS6 | HEXA | SLC7A11 | SHMT2 | PSMC1 |  |
|  | DEPTOR | KRAS | PFKM | ADK | COX4I1 | ARF5 |  |
| **Shared** | IDH3B | SLC3A2 | ALDH2 | LAMC1 | PGK1 | NAGLU |  |
| **Pam3CSK4** | SDHB | HLA-C | GAPDH | MSH2 | ITGB1 | MAP1LC3B |  |

### **Supplementary Table 1. Differentially expressed gene lists (padj<0.05, |FC|>1.25).**

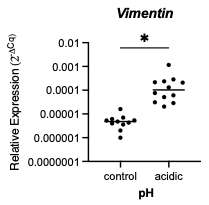

### **Supplementary Figure 2. Acid stress induces transcriptional upregulation of vimentin.**
